# Genomic architecture of human neuroanatomical diversity

**DOI:** 10.1101/001198

**Authors:** Roberto Toro, Jean-Baptiste Poline, Guillaume Huguet, Eva Loth, Vincent Frouin, Tobias Banaschewski, Gareth J Barker, Arun Bokde, Christian Büchel, Fabiana M Carvalho, Patricia Conrod, Mira Fauth-Bühler, Herta Flor, Jürgen Gallinat, Hugh Garavan, Penny Gowland, Andreas Heinz, Bernd Ittermann, Claire Lawrence, Hervé Lemaître, Karl Mann, Frauke Nees, Tomáš Paus, Zdenka Pausova, Marcella Rietschel, Trevor Robbins, Michael N Smolka, Andreas Ströhle, Gunter Schumann, Thomas Bourgeron, and the IMAGEN consortium (www.imagen-europe.com)^1^

**Author notes:** Corresponding author: Roberto Toro. Shared last authorship (Gunter Schumann, Thomas Bourgeron).

## Abstract

Human brain anatomy is strikingly diverse and highly inheritable: genetic factors may explain up to 80% of its variability. Prior studies have tried to detect genetic variants with a large effect on neuroanatomical diversity, but those currently identified account for <5% of the variance. Here we show, based on our analyses of neuroimaging and whole-genome genotyping data from 1,765 subjects, that up to 54% of this heritability is captured by large numbers of single nucleotide polymorphisms of small effect spread throughout the genome, especially within genes and close regulatory regions. The genetic bases of neuroanatomical diversity appear to be relatively independent of those of body size (height), but shared with those of verbal intelligence scores. The study of this genomic architecture should help us better understand brain evolution and disease.

## INTRODUCTION

Family studies show that a large part of the variability of different human brain structures is determined by genetic factors. Because we know a priori the degree of genetic relationship between monozygotic and dizygotic twins, or between members of a family, we can decompose the variability of a phenotype into genetic and environmental components. Various studies have demonstrated in this way that neuroanatomical phenotypes, such as brain volume or cortical surface, are highly inheritable, with genetic factors accounting for up to 80% of their variability (Winkler et al., 2010; Stein et al., 2012; Blokland, de Zubicaray, McMahon, & Wright, 2012). These results are particularly important for psychiatric research. Different psychiatric disorders have been associated with characteristic changes in brain anatomy, such as a higher incidence of macrocephaly and increases of white matter volume in autism (Amaral, Schumann, & Nordahl, 2008), or reduced hippocampal and total brain volumes in schizophrenia (Steen et al., 2012). If these characteristic changes are modulated by the subject’s genetic background, then this background may act as a protective factor or as a risk factor for the development of psychiatric conditions.

Whereas family studies can inform us about the heritability of a trait, different approaches are required to determine the nature of the genetic factors involved. Various efforts have been made to go deeper into the genetics of neuroanatomical diversity through candidate-gene approaches or through agnostic, genome-wide association studies (Bis et al., 2012; Ikram et al., 2012; Stein et al., 2012). These approaches have provided important insights on the genetic bases of neuroanatomical diversity, however, for the moment they account for only a small proportion of the phenotypic variance.

Here we used a recently developed approach (Yang et al., 2010, 2011), were the combined effect of hundreds of thousands SNPs is considered in a single test – instead of the massive univariate testing approach of classic genome-wide association studies (GWAS). We studied a large cohort of 1,765 adolescents from the IMAGEN project (Schumann et al., 2010), for whom neuroimaging, whole-genome genotyping and behavioural data was available. As in twin and family studies, we estimated the amount of phenotypic variance explained by genetic relationships among subjects. By contrast, instead of using expected relationships based on pedigree, we used a genome wide average of the difference in genotyping at each single nucleotide polymorphism (SNP) between unrelated subjects. By using different sets of SNPs to compute genetic relationships, we were able to partition neuroanatomical variance into different SNP sets and investigate the genomic architecture of neuroanatomical diversity at a level of granularity intermediate between that of family studies and candidate-gene or genome-wide association studies. Finally, we used simulated phenotypes to estimate the minimum number of causal SNPs likely to produce our observed results.

## RESULTS AND DISCUSSION

Brain scans were obtained from a cohort of 2,089 adolescents (14.5 ± 0.4 years old, 51% females) from the IMAGEN project (http://imagen-europe.com) using magnetic resonance imaging in 8 European centres. We measured intracranial volume (ICV), total brain volume (BV), as well as the volume of the hippocampus (Hip), thalamus (Th), caudate nucleus (Ca), putamen (Pu), globus pallidus (Pa), amygdala (Amy) and nucleus accumbens (Acc) using validated automatic segmentation programs (Buckner et al., 2004; Cox, 1996; Jenkinson, Bannister, Brady, & Smith, 2002; Patenaude, Smith, Kennedy, & Jenkinson, 2011; Smith et al., 2004; Zhang, Brady, & Smith, 2001) (Figure 1a, Supplementary Figure 1, Supplementary Table 1). Individuals were whole-genome genotyped using Illumina 610Quad and Illumina 660W-Quad chips. After various quality control filters, we conserved 269,308 informative, relatively independent (R^2^<0.9) SNPs in a cohort of 1,765 unrelated subjects.

**Figure 1.**
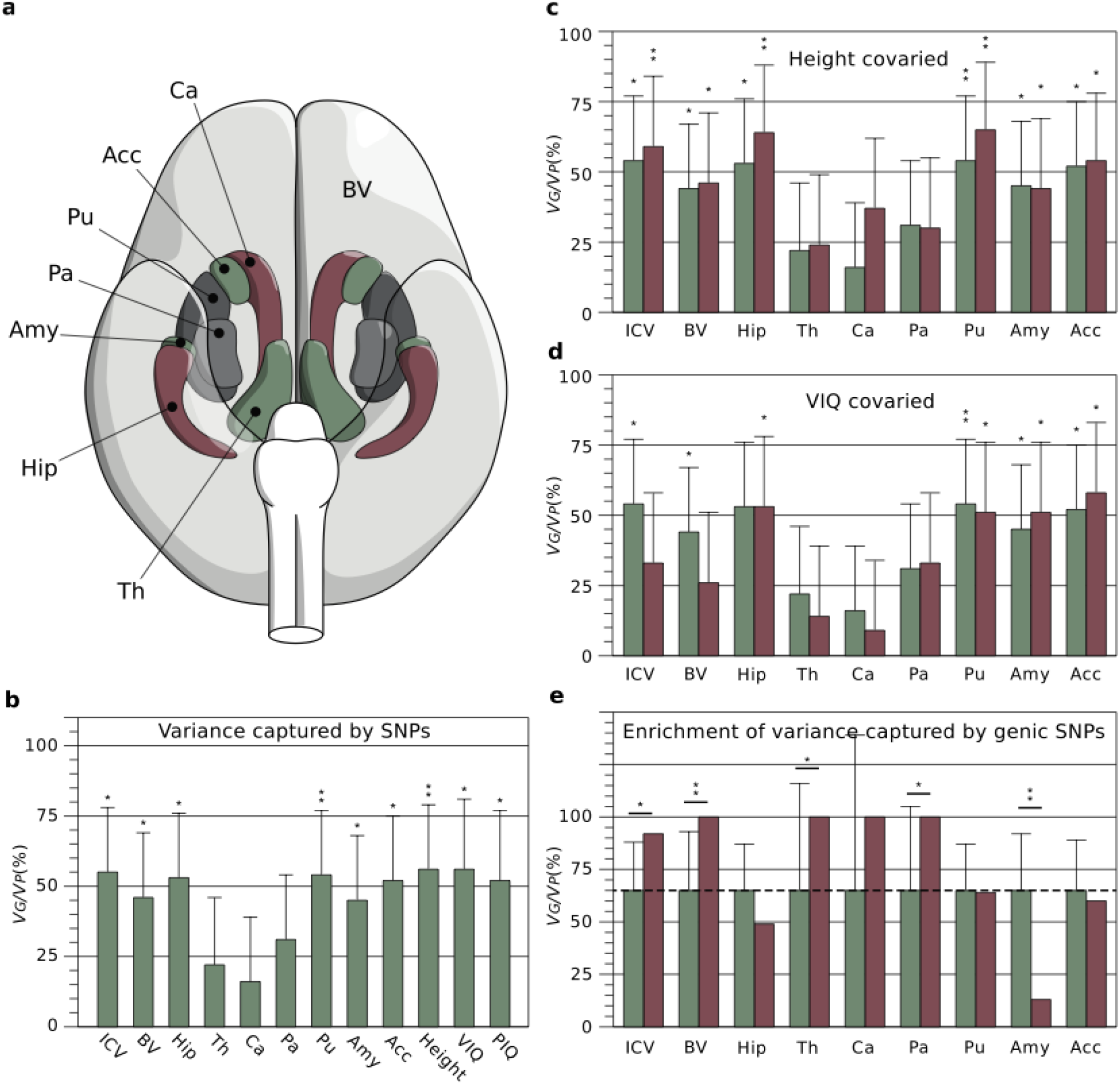
a. Brain phenotypes. We measured intracranial volume (not represented), total brain volume (BV, in light grey) and several subcortical structures, Ca: Caudate nucleus, Acc: Nucleus accumbens, Pu: Putamen, Pa: Pallidum, Amy: Amygdala, Hip: Hippocampus, and Th: Thalamus. **b. Variance captured by SNPs.** Percentage of phenotypic variance (*V_P_*) due to inter-individual genetic relationships (*V_G_*), computed from all genotyped SNPs. In addition to brain phenotypes, the bar plot includes estimates of *V_G_*/*V_P_* for height, VIQ and PIQ. **c. Effect of covarying body size (height) from brain phenotypes.** The proportion of *V_G_*/*V_P_* after covarying height (red bars) did not change substantially compared with those in 1b (green bars), and maintained their statistical significance. **d. Effect of covarying VIQ from brain phenotypes.** The proportion of *V_G_*/*V_P_* after covarying VIQ (red bars) decreased especially for ICV and BV, where the estimates were no longer statistically significant (green bars: raw estimates from 1b). **e. Enrichment of variance captured by genic SNPs.** Genic SNPs (gene boundaries ±50kbp) represent 64% of all SNPs. If all SNPs explained a similar amount of variance, genic SNPs should explain 64% of the total variance explained by SNPs (dashed line, green bars). They explained significantly more variance than expected for ICV, BV, Th and Pa; significantly less for Amy (red bars, error bars represent test variance). * P<0.05, ** P<0.01, uncorrected.

**Table 1.** Order of magnitude of the smallest P-value in the GWAS for each phenotype.

| Phenotype | $-\log_{10}(\text{Smallest P-value})$ |
| --- | --- |
| ICV | 5.95 |
| BV | 6.75 |
| Hip | 5.50 |
| Th | 5.72 |
| Ca | 5.91 |
| Pu | 5.49 |
| Pa | 5.67 |
| Amy | 5.14 |
| Acc | 5.60 |
| Height | 5.94 |
| VIQ | 5.32 |
| PIQ | 5.49 |

First, we estimated the proportion of the phenotypic variance explained by all SNPs with a linear mixed-effects model with the genetic relationship matrix as the structure of the covariance between subjects using GCTA (Yang et al., 2010). We es timated through simulation that we had >50% statistical power to find heritability values >45%, and >70% statistical power to find heritability values >55% (see Methods section). In all our analyses we included age, sex, and scanning centre as fixed effects. To account for population structure effects, we also in-cluded the first 10 principal components (PCs) of the identity-by-state (IBS) matrix as covariates (A. L. Price et al., 2006). Figure 1b shows the estimated proportion of the phenotypic variance attributable to genetic variance (*V_G_/V_P_*) for the neuroanatomical structures under study (Supplementary Table 2). The figure includes also estimates of *V_G_/V_P_* for height, as well as measurements of verbal intellectual quotient (VIQ) and performance intellectual quotient (PIQ) based on the Wechsler Intelligence Scale for Children. Our estimates for height (*V_G_/V_P_* = 56%, P = 0.0069), VIQ (*V_G_/V_P_* = 56%, P = 0.013) and PIQ (*V_G_/V_P_* = 52%, P = 0.02) were statistically significant, and consistent with those obtained previously in larger populations (Davies et al., 2011; Yang et al., 2010). A total of 12 statistical tests were performed. Because of the correlation among phenotypes a simple Bonferroni correction would be too conservative. Indeed, a global test shows that there is a statistically significant (P = 0.0011) excess of P-values <0.05 (Methods section).

We found that a large proportion of the variance of neuroanatomical phenotypes was explained by the additive effect of genotyped SNPs — for example, 44% (P = 0.031) of the variance in total brain volume (BV), 53% (P = 0.01) of the variance in hippocampal volume (Hip) and 54% (P = 0.011) of the variance in intracranial volume (ICV). Combined, the largest genome-wide association studies to date for BV (Stein et al., 2012), Hip (Bis et al., 2012; Stein et al., 2012) and ICV (Ikram et al., 2012; Stein et al., 2012) (N∼20,000) found 1 SNP associated to Hip volume and another associated to ICV, each explaining < 0.5% of the variance – of the same order of magnitude as for other quantitative traits, such as height. Approximately 50% of the additive genetic factors affecting neuroanatomical variability may be then supported by a large number of SNPs, each of small effect.

To get an idea of the minimum number of SNPs likely to produce our observed results we simulated 10,000 phenotypes with additive heritability of 50% produced by 1 to 1,000 causal SNPs and 10,000 phenotypes produced by 1 to 10,000 SNPs. Causal SNPs were randomly selected from among the original 518k genotyped SNPs before R^2^ filtering (i.e., their effect may be noticeable only through linkage disequilibrium), and their effect sizes drawn from a normal distribution to obtain 50% heritability. We performed GWAS for all our phenotypes, and recorded the order of magnitude of the smallest P-value, which varied from 10^−5.1^ to 10^−6.8^ (Table 1). We then did the same for each of the 20,000 simulated phenotypes. Figure 2 shows the proportion of simulations with smallest P-value of the order of 10^−5^, 10^−6^, etc., as a function of the number of causal SNPs used. We observed that 95% of the phenotypes simulated with <220 causal SNPs had P-values <10^−8^. By contrast, the order of the smallest P-value in the GWAS for ICV, for example, was 10^−5.95^, and 95% of simulations with smallest P-value <10^−5.95^ were produced by <850 causal SNPs. Similarly, the order of the smallest P-value in the GWAS for BV was 10^−6.8^, and 95% of the simulations with P-values smaller than that were produced by <420 causal SNPs. If the distribution of effect sizes of causal SNPs for ICV and BV were similar that used in our simulations, our phenotypes should likely be produced by hundreds of causal SNPs and possibly thousands of them.

**Figure 2.**
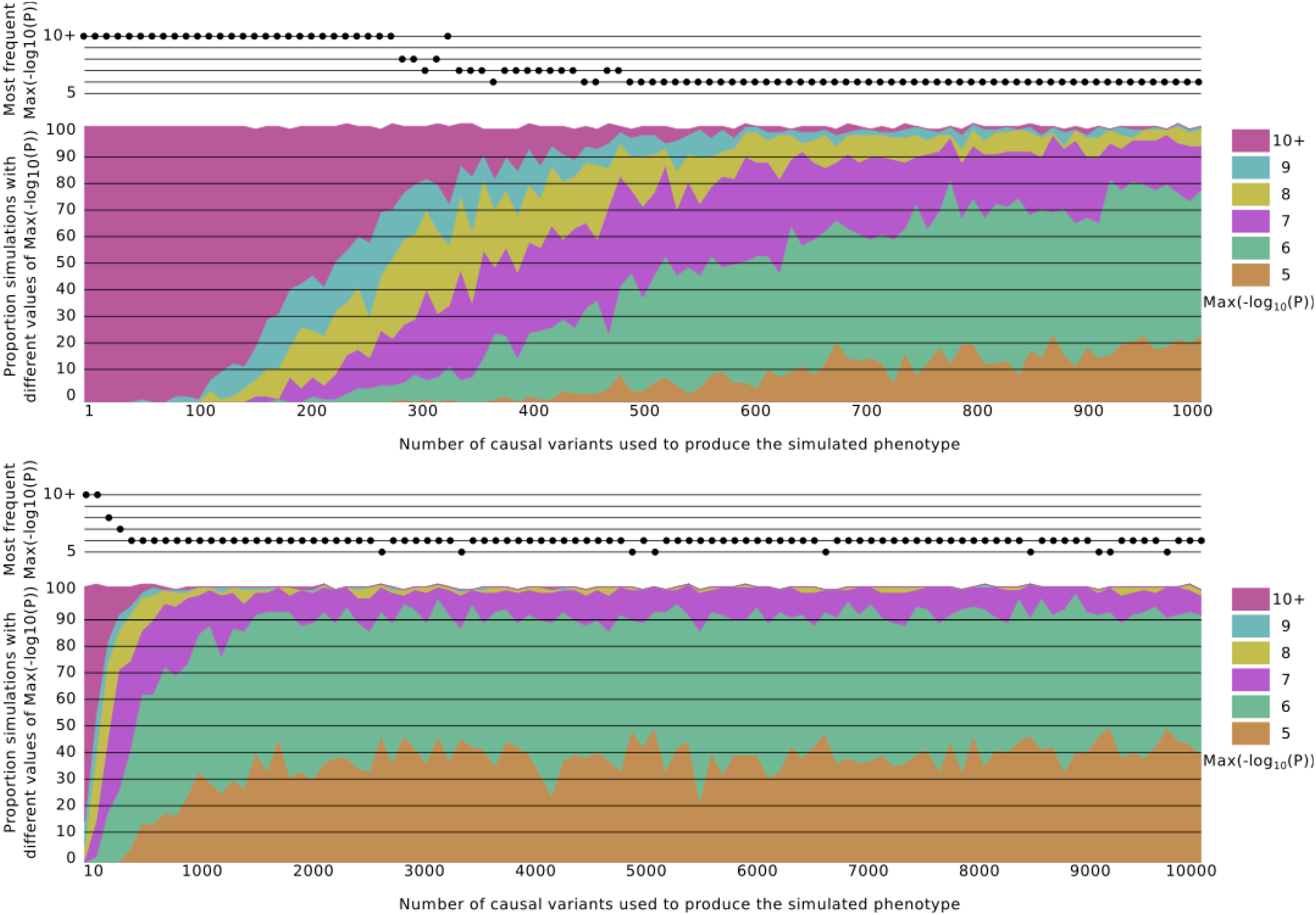
Distribution of smallest P-value in the GWAS with simulated phenotypes as a function of the number of causal SNPs used to generate them. Simulated phenotypes were produced with a number of causal SNPs varying from 1 to 1,000 (a), and from 1 to 10,000 (b). The effect of causal SNPs were drawn from a normal distribution, and the heritability of the simulated phenotypes was fixed at 50%. Ninety five percent of simulated phenotypes with <220 causal SNPs had a smallest P-value <10^−8^. By contrast, simulated phenotypes produced with >500 causal SNPs had most often a smallest P-value of the order of 10^−6^ or 10^−5^. The top plot in (a) and (b) shows the most frequent order of magnitude of the smallest P-value as a function of the number of causal SNPs.

The variance estimates for different brain structures were heterogeneous, and appeared to be differently related to height, VIQ and PIQ (Supplementary Table 2). For example, whereas the variance explained by SNPs was high and statistically significant for the hippocampus (*V_G_/V_P_* = 53%, P = 0.01), this was not the case for the caudate nucleus (*V_G_/V_P_* = 16%, P = 0.25) — a structure of comparable volume, geometry, and variability, that presents a similar correlation with ICV (r_Hip/ICV_ = 0.51, r_Ca/ICV_ = 0.52) and body size (r_Hip/Height_ = 0.15, r_Ca/Height_ = 0.21). This shows that the estimates of *V_G_/V_P_* are not merely determined by the structure’s volume or shape, and could rather reflect a varying influence of genetic and environmental factors. Our variance estimates were not significantly affected by population structure – not including the 10 first principal components of the IBS matrix changed on average the estimates of variance by less than 1% (P = 0.93). The estimates of variance did not change significantly either if height or PIQ were covaried (Figure 1c, Supplementary Table 2). By contrast, including VIQ scores as covariate decreased substantially *V_G_/V_P_* estimates for ICV and BV, but not for subcortical structures (Figure 1d). For example, ICV has a moderate correlation with height and VIQ (in our cohort r_ICV/Height_ = 0.39 and r_ICV/VIQ_ = 0.18). The estimate of *V_G_/V_P_* for ICV was not significantly different if height was added as a covariate, however, it decreased from 54% to 32% (no longer statistically significant) if VIQ was included as a covariate. We performed bivariate analyses to estimate the genetic correlation between our phenotypes, i.e., the amount of genetic variance shared by each pair of phenotypes (Supplementary Table 3). In particular, these analyses showed indeed a strong genetic correlation between VIQ and ICV (rG = 0.95, P = 0.0047), and between VIQ and BV (rG = 0.89, P = 0.014), but a small, not statistically significant, genetic correlation between height and ICV (rG = 0.20, P = 0.25), and between height and BV (rG = 0.23, P = 0.24). Genetic correlation was also weak between PIQ and ICV (rG = 0.02, P = 0.48) and between PIQ and BV (rG = 0.02, P = 0.48). More than 90% of brain volume (BV) is constituted by the cerebral cortex and its corticocortical connections. Our results suggest that the genetic bases of ICV and BV diversity may be shared to a larger extent with those of VIQ than with those of PIQ or body size (height).

A large proportion of the genetic variance captured by SNPs could be due to those located within genes and close regulatory regions. We obtained 20,022 gene boundaries from the UCSC Genome Browser hg18 assembly. We made a first set with all SNPs within these boundaries, and two further sets that included also SNPs 20kbp and 50kbp upstream and downstream from the 5’ and 3’ untranslated regions of each gene. Next, we computed genetic relationship matrices for those 3 SNP sets (±0kbp, ±20kbp and ± 50kbp genic sets), and their complements. Finally, for each of the 3 sets, we fitted the same linear mixed-effects model as before (including age, sex, centre and 10 principal components), but using 2 genetic relationship matrices instead of 1: the genic matrix and its complementary nongenic matrix. Genic SNP sets explained up to 98% of the variance captured by all SNPs (Supplementary Table 4), which was in many cases significantly larger than what could be expected from set length alone (Figure 1e, Supplementary Table 5). For ICV, where 54% of the variance can be explained by all genotyped SNPs (N = 273,926), using only SNPs within gene boundaries (N = 108,339) explained 26% of the phenotypic variance (P = 0.054), and this proportion increased to 45% (P = 0.0065) when the boundaries were expanded to ± 20kbp (N = 146,431), and to 49% (P = 0.0058) when the boundaries were expanded to ± 50kbp (N = 174,334). The genic ± 50kbp set contained 64% of all genotyped SNPs, but explained 91% of the variance of ICV attributable to SNPs, significantly more than what we would expect from its length alone (P = 0.014).

Previous reports have suggested that causal SNPs for height and IQ are relatively homogeneously distributed across the genome, and then, that increasing the number of SNPs used to create a genetic-relationship matrix increases proportionally the amount of phenotypic variance captured. We observed the same trend for our neuroanatomical phenotypes. We partitioned the genome into non-overlapping sets with different numbers of SNPs, and observed a strong correlation between set length and *V_G_/V_P_* (r = 0.62 on average). The correlation was the same when only genic SNPs were selected (r = 0.62), but smaller, and in most cases not statistically significant when only nongenic SNPs were selected (Figure 3).

**Figure 3.**
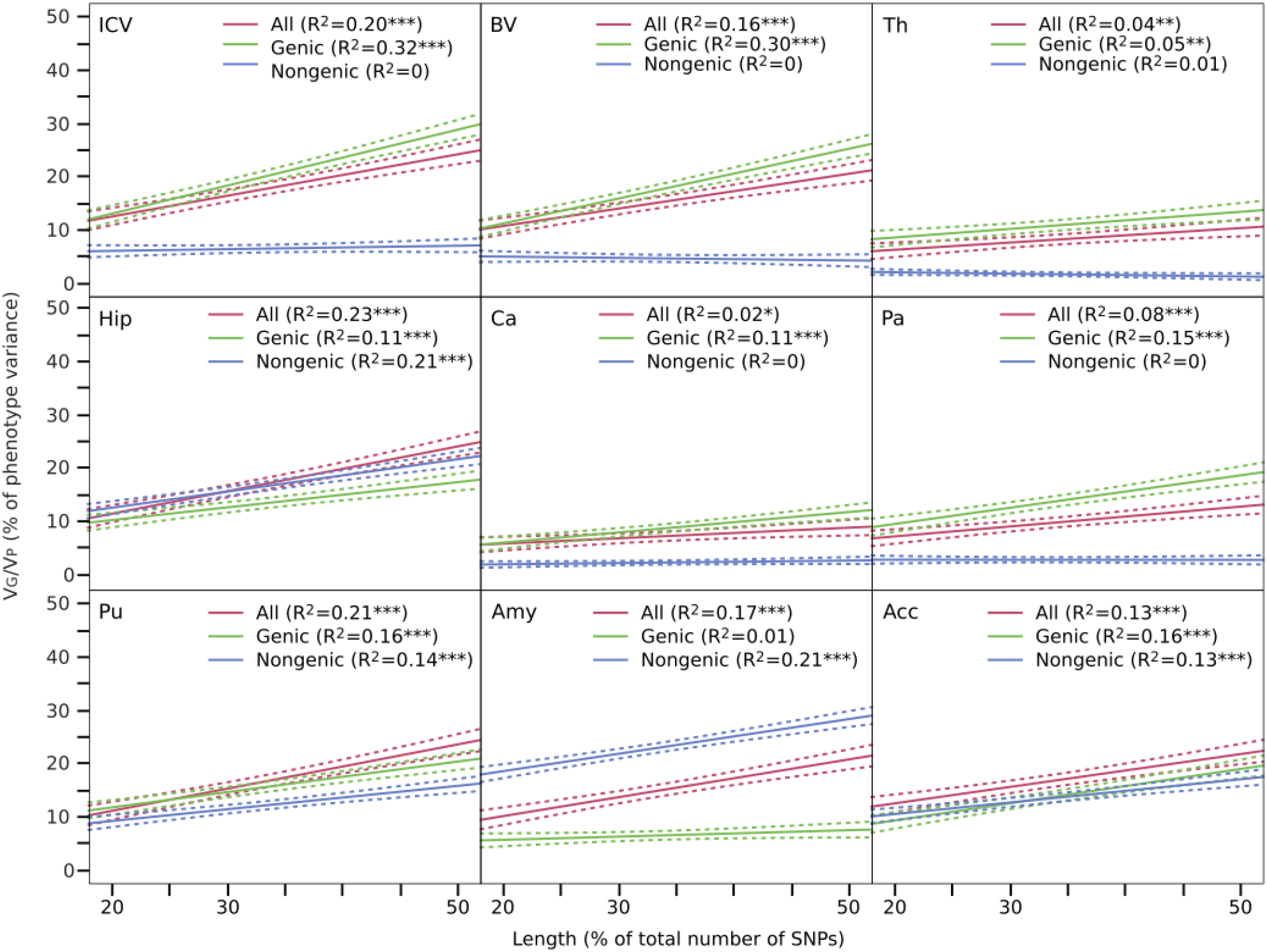
*V_G_*/*V_P_* versus gene set length. The amount of variance captured by SNPs increased with the number of SNPs used to compute genetic-relationship matrices. In most cases, this was only the case for genic SNPs (Ref.Seq.±50kpb). * P<0.01, ** P<0.001, *** P<0.0001, uncorrected.

Finally, we partitioned *V_G_/V_P_* based on functional annotation (SNPs within genes involved in central nervous system function (Lee et al., 2012; Raychaudhuri, Korn, & McCarroll, 2010)), and relative minor-allele frequency. We did not observe statistically significant differences in the amount of variance explained by these different SNP sets compared with the expectations based on their length (Supplementary Methods, Supplementary Tables 6, 7).

In conclusion, our analyses indicate that a significant proportion of the heritability of neuroanatomical phenotypes may result from the additive effect of hundreds of small-effect SNPs spread genome-wide. Such SNPs seemed to be largely independent from those related to body size (height) or reflecting population structure in our cohort. They were shared to a greater extent, however, with those associated with VIQ in the case of ICV and BV. An especially important role in determining neuroanatomical diversity appeared to be played by SNPs within genes and close regulatory regions.

Even if our variance estimates are large, they are still far from the estimates of additive genetic variance from pedigree studies: ∼80% of the variance of various brain structures has been attributed to additive genetic factors (Stein et al., 2012). This difference may be due to a weak linkage disequilibrium between the genotyped SNPs and the real causative variants, to rare alleles with larger effect sizes or to common alleles with even smaller effect sizes. In any case, further progress will require cohorts of maybe hundreds of thousands of individuals, underlining the necessity for international efforts such as the ENIGMA and CHARGE consortia.

Recent studies have highlighted the importance of the additive effect of SNPs in determining anatomical and cognitive diversity in humans, but also their role in psychiatric disorders. In addition to the clear role of rare mutations in the susceptibility to psychiatric disorders (Cook Jr & Scherer, 2008), whole-genome analyses of variance have shown that commonly genotyped SNPs capture 23% of the risk to schizophrenia (CrossDisorder Group of the Psychiatric Genomics Consortium, 2013; Lee et al., 2012), 24% of the risk to Alzheimer’s disease (Lee et al., 2013), and from 17% to 60% of the risk to autism spectrum disorders (Cross-Disorder Group of the Psychiatric Genomics Consortium, 2013; Klei et al., 2012). Due to the small individual effect of these SNPs, GWAS will require very large cohorts to explain any sizeable proportion of the trait’s genetic variance (Park et al., 2010). Various structural and functional neuroimaging endophenotypes, on the other hand, have been frequently associated with psychiatric disorders (Meyer-Lindenberg & Weinberger, 2006), and their analysis using whole-genome regression could inform us about the added effect of SNPs at a relevant intermediate level, closer to biological processes than cognitive or psychiatric tests. A global view of the genomic architecture of neuroimaging endophenotypes should not only allow us to better understand the biological bases of the susceptibility to psychiatric disorders – helping us, for example, to target future GWAS to more specific chromosomal regions and brain structures – but also to improve our understanding of the biological bases of brain diversity and evolution in humans.

## METHODS

### Neuroimaging

Magnetic resonance imaging data were acquired at 8 European centres, using a standardised 3 Tesla, T1-weighted gradient echo protocol (voxel size = 1.1 mm isotropic) based on that from the ADNI initiative (http://adni.loni.ucla.edu/methods/documents/mri-protocols). MRI volumes were first linearly transformed to match the MNI152 atlas using FLIRT from FSL (Jenkinson et al., 2002; Smith et al., 2004) (http://fsl.fmrib.ox.ac.uk/fsl/fslwiki/FLIRT). The inverse of the determinant of the transformation matrix was used to estimate intracranial volume (Buckner et al., 2004). Next, skull was stripped using 3dSkullStrip from AFNI (Cox, 1996) (http://afni.nimh.nih.gov), and the grey matter, white matter and cerebrospinal fluid were automatically segmented using FAST (Zhang et al., 2001) (http://fsl.fmrib.ox.ac.uk/fsl/fslwiki/FAST). The skull-stripped versions of the brain volumes, and the tissue segmentations were visually inspected and manually corrected wherever necessary. Total brain volume was estimated as the sum of total grey and white matter volumes. Finally, subcortical structures were automatically segmented using FIRST (Patenaude et al., 2011) (http://fsl.fmrib.ox.ac.uk/fsl/fslwiki/FIRST), and their accuracy visually controlled using in-house software. All volumes were log10 converted. Their distribution is illustrated in Fig. S1. Despite the differences in average volume (from ∼1 cm^3^ for the amygdala, to ∼1,300 cm^3^ for total brain volume, Fig. S1), all structures showed a similar variability – there was a ∼1.8-fold change from the smallest to the largest volume in the cohort. The correlation matrix for all phenotypes analysed is shown in Table S1.

### Genotyping and quality control

We used the autosomal SNPs common to the Illumina 610-Quad and Illumina 660W-Quad chips, and strict filtering to conserve high-quality SNPs only (minor-allele frequency >5%, genotyping rate >99%, significance threshold for Hardy-Weinberg equilibrium test <10^−6^, subjects missing genotyping <10%, using PLINK (Purcell et al., 2007), http://pngu.mgh.harvard.edu/∼purcell/plink). We further excluded SNPs in strong linkage disequilibrium (R^2^ > 0.9) within a window of 50 SNPs to prevent colinearity in our analyses. The final genotyping data consisted of 269,308 SNPs.

### Estimation of variance captured by SNPs

The variance of a phenotype attributable to genetic factors is classically estimated by comparing the correlation between pairs of monozygotic (MZ) and dizygotic (DZ) twins. Pairs of MZ and DZ twins share a common maternal environment during foetal life, but MZ twins share 100% of their genomes whereas DZ twins share on average 50% of it. If the variance of the phenotype is affected by genetic factors, the correlations between MZ twins will be larger than those observed in DZ twins. The amount of variance due to genetic factors can be then estimated using Falconer’s formula (Falconer, 1965), or more accurately, by using the restricted maximum likelihood method (Corbeil & Searle, 1976) (REML). The twin study design can be extended to the analysis of more complex family relationships, by using the levels of relationship expected from the pedigrees. In neuroscience, this approach has been successfully used to show that genetic factors explain an important part of the variance of several neuroanatomical phenotypes such as brain volume (Stein et al., 2012), cortical surface extension (Winkler et al., 2010) or white matter microstructure (Kochunov et al., 2010).

Twin and extended pedigrees studies provide important information on the role of genetic factors in determining neuroanatomical phenotypes, but complementary approaches are needed to investigate the nature of these genetic factors. In the recent years, several research groups have attempted to discover the genetic bases of the high heritability of neuroanatomical phenotypes by studying, for example, their association with SNPs in candidate genes (BDNF (Pezawas et al., 2004), microcephaly genes (Rimol et al., 2010), etc.) or through genome-wide association studies (Bis et al., 2012; Ikram et al., 2012; Stein et al., 2012). Yet these findings explain today only a minute part of the phenotypic variability.

The approach that we used here, implemented by the GCTA software (Yang et al., 2010, 2011) (http://www.complextraitgenomics.com/software/gcta), estimates the variance captured by a large number of SNPs (modelled as random effects), providing information at a level intermediate between twin studies and association studies. Like twin and extended pedigree studies, GCTA estimates the part of the phenotypic variance due to the matrix of genetic relationships among subjects. But instead of using levels of genetic relationship expected a priori from a pedigree, these levels are computed from genotyping data (Lynch & Ritland, 1999; Ritland, 1996; Yang et al., 2010). The relationship between phenotype variance *Var(y)*, the variance of the additive genetic effects 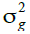, and the residual variance 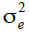 is formulated as follows

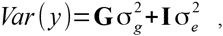

where **G** is the genetic relationship matrix, containing the degree of genetic relationship, and **I** is the identity matrix. In GCTA the level of genetic relationship between each pair of individuals *j* and *k* – the entries of the **G** matrix – is calculated as a weighted average across all SNPs:

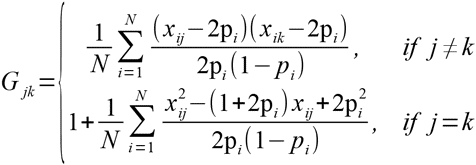

where *x_ij_* equals 0, 1 or 2 depending on whether the genotype of the *i*-th SNP of the *j*-th subjects is AA, AB or BB; *p*_*i*_ is the allele frequency of SNP *i*; and *N* is the number of SNPs considered in the analysis (in our case, *N* = 269,308). The level of genetic relationship between two subjects is then a single value summarising how similar their genomes are.

GCTA estimates variance components using the Average Information REML (Gilmour, Thompson, & Cullis, 1995) (AI-REML) method – a variant of the classic REML that provides a more efficient estimation even if the **G** matrix is large. The statistical significance of the genetic variance estimates (the P-values reported here) were computed using a maximum likelihood ratio test (LRT) comparing the complete model which includes the genetic effect, to a partial model which excludes it. In REML analyses, LRT values are distributed as a 50% mixture of 0 and a Chi-square with degrees of freedom (df) equal to the number of genetic relationship matrices being tested. The LRT values and the corresponding degrees of freedom for the tests are indicated in the LRT column of Supplementary Tables S2, S4, S6-7.

### Confounding factors

Several confounding factors could affect our variance estimates. For all our analyses we included age, sex and scanning centre as covariates. Additional analyses including Pubertal Development Scale scores (Carskadon & Acebo, 1993) did not affect the results and this covariate was no longer included in the model. Population structure might also affect neuroanatomical diversity or bias our variance estimates:

1. Our estimates could be affected by cryptic relatedness if our cohort included subjects distantly related. In that case, phenotypic similarity could be partly due to shared environment effects or familial causal variants not captured by SNPs. To prevent this, we excluded subjects with a genetic relationship > 0.025 (i.e., more related than 3rd or 4th cousins).
2. We used Admixture (Alexander, Novembre, & Lange, 2009) (http://www.genetics.ucla.edu/software/admixture) to estimate individual ancestry relative to the reference populations in HapMap 3 (“The International HapMap Project.,” 2003) (http://hapmap.ncbi.nlm.nih.gov). The result (Fig. S2) showed that individuals in our cohort have a strong European ancestry component.
3. It has been shown that including as covariates the first principal components of the matrix of identity-by-state (IBS) distance between subjects efficiently accounts for population structure effects (A. L. Price et al., 2006; A. Price, Zaitlen, Reich, & Patterson, 2010). Usually, the first 4 or 5 principal components are included. Here, we included the first 10 principal components in all our analyses. We observed, however, that not including them affected only marginally our variance estimates. On average, the difference between estimates including and excluding the first 10 principal components of the IBS matrix was of 0.6%, not significantly different (2tailed t-test, P = 0.9346).
4. Partitioning the variance explained by SNPs between genic and nongenic showed that in many cases the former explained significantly more variance than the latter (see bellow for a description of the method). For example, SNPs within genes ± 50kbp explain 91% of the total variance of ICV captured by SNPs. If our estimates were driven by population structure effects, we could expect an excess of Ancestry Informative Markers (AIMs) within the genic SNP set. We obtained a list of 1,442 AIMs from Tian and collaborators (Tian et al., 2008), 604 of which were contained in our SNP list. There was no statistically significant difference in the number of AIMs between our genic and nongenic SNP sets (375 genic AIM versus 229 nongenic AIMs, Fisher’s exact test P = 0.1723), and if anything, there was a tendency for AIMs to be underrepresented within the genic SNP set (Fisher’s exact test P = 0.0892).

These analyses suggest that population stratification effects did not play a major role in the determination of neuroanatomical variability in our cohort (Table S2).

### Estimation of statistical power

We simulated 10,000 phenotypes with different heritability values, supported by a different number of causal SNPs. We uniformly sampled heritability values in the range from 0 to 80%, and number of causal SNPs from 1 to 10,000. The causal variants were selected from the non-pruned list of SNPs (∼518k SNPs), but the genetic relationship matrices were computed using only SNPs from the pruned set (∼270k SNPs). In consequence, the effect of some of the causal variants would be only captured through their linkage disequilibrium with the SNPs retained in the pruned list. Statistical power achieved to detect a given heritability was estimated as the proportion of test with P < 0.05 (Figure 4).

**Figure 4.**
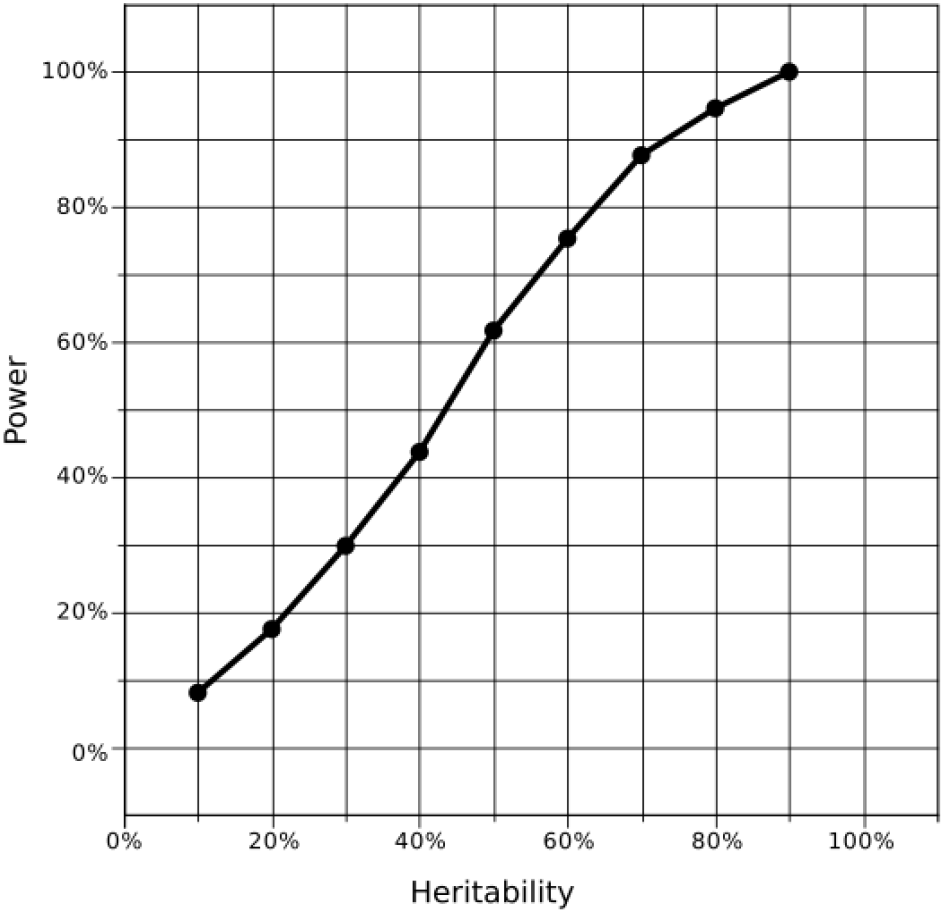
Statistical power as a function of heritability. Estimation of statistical power obtained through simulation of 10,000 phenotypes with different heritability values, supported by a different number of causal SNPs. We had >50% statistical power to find heritability values >45%, and >70% statistical power to find heritability values >55%.

### Correlation between SNP set size and V_G_/V_P_

We constructed genetic relationship matrices for 3 sets of nonoverlapping, randomly selected, SNPs of small, medium and large size. These sets were drawn from all genotyped SNPs, or only from genic SNPs (Ref. Seq. ± 50kbp), or nongenic SNPs. We ensured that small, medium and large sets contained the same number of SNPs in all 3 groups by selecting 20%, 30% and 50% of the total number of nongenic SNPs, the less numerous group. We performed 100 repetitions of this procedure, each time randomly selecting non-overlapping sets of 20%, 30% and 50% (20%+30%+50%=100%) of SNPs from all genotyped SNPs, or only from the genic or nongenic subgroups. For each repetition, we computed the correlation between *V_G_/V_P_* and set size. Correlation coefficients were converted to Z values using Fisher’s transformation, and the distribution tested against the null-hypothesis of no correlation (2-tailed t-test). The amount of variance of ICV, BV, subcortical volumes, height, VIQ and PIQ explained by the low, medium and long sets correlated significantly with the size of the SNP set (Figure 2).

### Partition of V_G_/V_P_

We partitioned *V_G_/V_P_* among non-overlapping sets of SNPs, for example, genic and nongenic SNPs (2 sets) or SNPs of low, medium and high minor-allele frequency (MAF, 3 sets), etc. We computed a genetic relationship matrix **G_i_** for each of these *n* sets, and used them as random effects in our model. The variance of our phenotypes *Var* (*y*) was therefore decomposed as

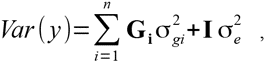

where the number of sets would be n = 2 for the case of a genic versus nongenic partition, or n = 3 in the case of a partition into low, medium and high MAF.

The LRT columns in Supplementary Tables S4, S6-7 indicate the value of the likelihood ratio test comparing the complete model (including all genetic relationship matrices) to the partial model that includes only the residual variance component. The Pmodel columns indicate the statistical significance of these values given the number of variance components tested (degrees of freedom).

As *a posteriori* analyses, we tested whether the variance explained by one of these sets, genic SNPs for example, was larger than what could be expected given its number of SNPs. The total genetic variance explained is

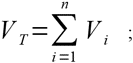

where *N* is the total number of SNPs, and *N_i_* the number of SNPs in set *i*, *i* = 1,…,*n*. If all SNP sets were equivalent, then the amount of variance they explain should be simply proportional to their length, and then

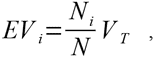

where *EV_i_* is the expected amount of variance explained by the *i*-th set. We wanted to test whether the difference *V_i_-EV_i_* was significantly larger than 0, so we constructed a *Z*-score

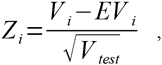

where

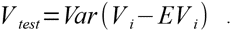

Note that *V_i_* here is the estimated explained variance for group *i* – a random variable; whereas *EV_i_* is a fixed value. We compared the observed value of *Z_i_* with those obtained from >10,000 random permutations, where *n* non-overlapping SNP sets of size *N_i_* were randomly sampled from all available SNPs (without replacement).

### Partition of V_G_/V_P_ based on involvement in central nervous system function

We looked at the proportion of *V_G_/V_P_* that could be attributed to genes preferentially expressed in the central nervous system, playing a role in neuronal activity, learning, or involved in synaptic function. We used the set of 2,725 genes defined by Raychaudhuri and collaborators (Raychaudhuri et al., 2010) and previously used in the SNP-based heritability analyses of the susceptibility to schizophrenia by Lee and collaborators (Lee et al., 2012). We made 3 SNP sets: the 1st set, CNS+, contained all SNPs within ± 50 kbp of the 5’ and 3’ UTR of the gene set (N = 61,175, 23% of the total number of SNPs); the 2nd set, CNS-, contained all the remaining genic SNPs (N = 113,160, 42% of the total number of SNPs); and the 3rd set regrouped all nongenic SNPs. As before, the genetic-relationship matrices computed using these 3 SNP sets were used in a single linear mixed model. We found that the amount of variance explained by the CNS+ set was not significantly different than what we expected from its size (Table S6).

### Partition of V_G_/V_P_ based on MAF

Allele frequency variations may provide hints about the evolutionary history of a trait. We estimated the proportion of *V_G_/V_P_* that can be attributed to sets of SNPs with low (5-20%), medium (20-35%) and high (35-50%) minor allele frequencies. SNPs in the low-frequency set were the most numerous, 48% of all SNPs, followed by medium-frequency SNPs (30%), and high-frequency alleles (22%). Table S7 shows the result of fitting a linear mixed model with the 3 genetic-relationship matrices computed using the low, medium and high-frequency, in addition to the same fixed effects as previously. We could expect each set to explain a fraction of the variance corresponding to the proportion of the total number of SNPs they represent. Furthermore, because SNPs of high MAF are individually more informative than those with low minor-allele frequency, they could potentially explain more variance (the variance of the genetic-relationship matrices increased from the low to the medium to the high frequency set). However, the amount of variance explained by the different sets was not significantly larger than what we expected from their size.

### Global test on P-values

We performed our analyses on 12 correlated phenotypes (9 brain regions, plus Height, VIQ and PIQ). Because of these correlations, a standard Bonferroni correction would be too conservative. Indeed, after Bonferroni correction, just a few results would remain statistically significant. However, under the null hypothesis we should expect around 5% of these tests to be significant, but the observed number of P-values <0.05 was much larger. To test the significance of this excess we constructed a statistic S from the list of P-values converted to Z-values obtained for each phenotype:

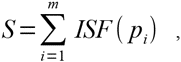

where *m = 12* is the number of tests performed and ISF stands for the inverse survival function of the normal distribution. We then generated the distribution of *S* under the null hypothesis by drawing from a multivariate Gaussian distribution with a variance-covariance structure given by the correlation matrix across phenotypes (Table S1). The significance of the excess of P-values was estimated as the proportion of scores under the null hypothesis that were greater than the observed score. The result of this global test is indicated at the final row of supplementary tables S2, S4-6.

## ACKNOWLEDGEMENTS

This work was supported by the European Union-funded FP6 Integrated Project IMAGEN (Reinforcement-related behaviour in normal brain function and psychopathology) (LSHM-CT-2007-037286), the FP7 projects ADAMS (Genomic variations underlying common neuropsychiatric diseases and diseases related to cognitive traits in different human populations) (242257), the Innovative Medicine Initiative Project EU-AIMS (115300-2), the Medical Research Council Programme Grant “Developmental pathways into adolescent substance abuse” (93558), the Swedish Funding Agency FORMAS, the German Bundesministerium und Forschung (FKZ: 01EV0711), Institut Pasteur, CNRS, Université Paris Diderot, the Betten-court-Schueller Foundation, the Conny-Maeva Foundation, the Orange Foundation, the FondaMental Foundation and the Cognacq-Jay Foundation. RT thanks Hong Lee, Jian Yang and Naomi Wray for their help.

## REFERENCES

Alexander, D. H., Novembre, J., & Lange, K. (2009). Fast model-based estimation of ancestry in unrelated individuals. Genome Research, 19(9), 1655–64. doi:10.1101/gr.094052.109

Amaral, D. G., Schumann, C. M., & Nordahl, C. W. (2008). Neuroanatomy of autism. Trends in neurosciences, 31(3), 137–45. doi:10.1016/j.tins.2007.12.005

Bis, J. C., DeCarli, C., Smith, A. V., van der Lijn, F., Crivello, F., Fornage, M., … Srikanth, V. (2012). Common variants at 12q14 and 12q24 are associated with hippocampal volume. Nature genetics, 44(5), 545–51. doi:10.1038/ng.2237

Blokland, G. A. M., de Zubicaray, G. I., McMahon, K. L., & Wright, M. J. (2012). Genetic and environmental influences on neuroimaging phenotypes: a meta-analytical perspective on twin imaging studies. Twin research and human genetics: the official journal of the International Society for Twin Studies, 15(3), 351–371. doi:10.1017/thg.2012.11

Buckner, R. L., Head, D., Parker, J., Fotenos, A. F., Marcus, D., Morris, J. C., & Snyder, A. Z. (2004). A unified approach for morphometric and functional data analysis in young, old, and demented adults using automated atlas-based head size normalization: reliability and validation against manual measurement of total intracranial volume. NeuroImage, 23(2), 724– 38. doi:10.1016/j.neuroimage.2004.06.018

Carskadon, M. A., & Acebo, C. (1993). A self-administered rating scale for pubertal development. The Journal of adolescent health: official publication of the Society for Adolescent Medicine, 14(3), 190–195.

Cook Jr, E. H., & Scherer, S. W. (2008). Copy-number variations associated with neuropsychiatric conditions. Nature, 455(7215), 919–923. doi:10.1038/nature07458

Corbeil, R. R., & Searle, S. R. (1976). Restricted maximum likelihood (REML) estimation of variance components in the mixed model. Technometrics, 18(1), 31–38.

Cox, R. W. (1996). AFNI: software for analysis and visualization of functional magnetic resonance neuroimages. Computers and biomedical research, 29(3), 162–73.

Cross-Disorder Group of the Psychiatric Genomics Consortium. (2013). Genetic relationship between five psychiatric disorders estimated from genome-wide SNPs. Nature Genetics, 45(9), 984–994. doi:10.1038/ng.2711

Davies, G., Tenesa, A., Payton, A., Yang, J., Harris, S. E., Liewald, D., … Deary, I. J. (2011). Genome-wide association studies establish that human intelligence is highly heritable and polygenic. Molecular psychiatry, 16(10), 996–1005. doi:10.1038/mp.2011.85

Falconer, D. (1965). The inheritance of liability to certain diseases, estimated from the incidence among relatives. Annals of Human Genetics, 29(1), 51–76. doi:10.1111/j.1469-1809.1965.tb00500.x

Gilmour, A. R., Thompson, R., & Cullis, B. R. (1995). Average Information REML: An Efficient Algorithm for Variance Parameter Estimation in Linear Mixed Models. Biometrics, 51(4), 1440–1450.

Ikram, M. A., Fornage, M., Smith, A. V., Seshadri, S., Schmidt, R., Debette, S., … Taal, H. R. (2012). Common variants at 6q22 and 17q21 are associated with intracranial volume. Nature genetics, 44(5), 539–44. doi:10.1038/ng.2245

Jenkinson, M., Bannister, P., Brady, M., & Smith, S. (2002). Improved optimization for the robust and accurate linear registration and motion correction of brain images. NeuroImage, 17(2), 825–841.

Klei, L., Sanders, S. J., Murtha, M. T., Hus, V., Lowe, J. K., Willsey, A. J., … Devlin, B. (2012). Common genetic variants, acting additively, are a major source of risk for autism. Molecular Autism, 3(1), 9–9. doi:10.1186/2040-2392-3-9

Kochunov, P., Glahn, D. C., Lancaster, J. L., Winkler, A. M., Smith, S., Thompson, P. M., … Blangero, J. (2010). Genetics of microstructure of cerebral white matter using diffusion tensor imaging. NeuroImage, 53(3), 1109–16. doi:10.1016/j.neuroimage.2010.01.078

Lee, S. H., DeCandia, T. R., Ripke, S., Yang, J., Sullivan, P. F., Goddard, M. E., … Wray, N. R. (2012). Estimating the proportion of variation in susceptibility to schizophrenia captured by common SNPs. Nature genetics, 44(3), 247–50. doi:10.1038/ng.1108

Lee, S. H., Harold, D., Nyholt, D. R., Goddard, M. E., Zondervan, K. T., Williams, J., … Visscher, P. M. (2013). Estimation and partitioning of polygenic variation captured by common SNPs for Alzheimer’s disease, multiple sclerosis and endometriosis. Human Molecular Genetics, 22(4), 832–41. doi:10.1093/hmg/dds491

Lynch, M., & Ritland, K. (1999). Estimation of pairwise relatedness with molecular markers. Genetics, 152(4), 1753– 66.

Meyer-Lindenberg, A., & Weinberger, D. R. (2006). Intermediate phenotypes and genetic mechanisms of psychiatric disorders. Nature Reviews Neuroscience, 7(10), 818–27. doi:10.1038/nrn1993

Park, J.-H., Wacholder, S., Gail, M. H., Peters, U., Jacobs, K. B., Chanock, S. J., & Chatterjee, N. (2010). Estimation of effect size distribution from genome-wide association studies and implications for future discoveries. Nature genetics, 42(7), 570–5. doi:10.1038/ng.610

Patenaude, B., Smith, S. M., Kennedy, D. N., & Jenkinson, M. (2011). A Bayesian model of shape and appearance for subcortical brain segmentation. NeuroImage, 56(3), 907–22. doi:10.1016/j.neuroimage.2011.02.046

Pezawas, L., Verchinski, B. A., Mattay, V. S., Callicott, J. H., Kolachana, B. S., Straub, R. E., … Weinberger, D. R. (2004). The brain-derived neurotrophic factor val66met polymorphism and variation in human cortical morphology. The Journal of Neuroscience, 24(45), 10099–102. doi:10.1523/JNEUROSCI.2680-04.2004

Price, A. L., Patterson, N. J., Plenge, R. M., Weinblatt, M. E., Shadick, N. A., & Reich, D. (2006). Principal components analysis corrects for stratification in genome-wide association studies. Nature genetics, 38(8), 904–9. doi:10.1038/ng1847

Price, A., Zaitlen, N., Reich, D., & Patterson, N. (2010). New approaches to population stratification in genome-wide association studies. Nature Reviews Genetics, 11(7), 459–63. doi:10.1038/nrg2813

Purcell, S., Neale, B., Todd-Brown, K., Thomas, L., Ferreira, M. A. R., Bender, D., … Sham, P. C. (2007). PLINK: a tool set for whole-genome association and population-based linkage analyses. American Journal of Human Genetics, 81(3), 559–75. doi:10.1086/519795

Raychaudhuri, S., Korn, J., & McCarroll, S. (2010). Accurately assessing the risk of schizophrenia conferred by rare copynumber variation affecting genes with brain function. PLoS Genetics, 6(9). doi:10.1371/journal.pgen.1001097

Rimol, L. M., Agartz, I., Djurovic, S., Brown, A. a, Roddey, J. C., Kähler, A. K., … Andreassen, O. a. (2010). Sex-dependent association of common variants of microcephaly genes with brain structure. Proceedings of the National Academy of Sciences of the United States of America, 107(1), 384–8. doi:10.1073/pnas.0908454107

Ritland, K. (1996). A Marker-Based Method for Inferences About Quantitative Inheritance in Natural Populations. Evolution, 50(3), 1062–1062. doi:10.2307/2410647

Schumann, G., Loth, E., Banaschewski, T., Barbot, A., Barker, G., Büchel, C., … Struve, M. (2010). The IMAGEN study: reinforcement-related behaviour in normal brain function and psychopathology. Molecular psychiatry, 15(12), 1128–39. doi:10.1038/mp.2010.4

Smith, S. M., Jenkinson, M., Woolrich, M. W., Beckmann, C. F., Behrens, T. E. J., Johansen-Berg, H., … Matthews, P. M. (2004). Advances in functional and structural MR image analysis and implementation as FSL. NeuroImage, 23 Suppl 1 (Supplement 1), S208–S219.

Steen, R. G., Mull, C., Mcclure, R., Hamer, R. M., Jeffrey, A., Steen, A. N. T., & Lieberman, J. A. (2012). Brain volume in first-episode schizophrenia : Systematic review and metaanalysis of magnetic resonance imaging studies REVIEW ARTICLE AUTHOR ’S PROOF Brain volume in first-episode schizophrenia Systematic review and meta-analysis of magnetic resonance i, i 510–518. doi:10.1192/bjp.188.6.510

Stein, J. L., Medland, S. E., Vasquez, A. A., Hibar, D. P., Senstad, R. E., Winkler, A. M., … Bergmann, Ø. (2012). Identification of common variants associated with human hippocampal and intracranial volumes. Nature genetics, 44(5), 552–61. doi:10.1038/ng.2250

The International HapMap Project. (2003). Nature, 426(6968), 789–96. doi:10.1038/nature02168

Tian, C., Plenge, R. M., Ransom, M., Lee, A., Villoslada, P., Selmi, C., … Seldin, M. F. (2008). Analysis and application of European genetic substructure using 300 K SNP information. PLoS Genetics, 4(1), e4–e4. doi:10.1371/journal.pgen.0040004

Winkler, A. M., Kochunov, P., Blangero, J., Almasy, L., Zilles, K., Fox, P. T., … Glahn, D. C. (2010). Cortical thickness or grey matter volume? The importance of selecting the phenotype for imaging genetics studies. NeuroImage, 53(3), 1135– 46. doi:10.1016/j.neuroimage.2009.12.028

Yang, J., Benyamin, B., McEvoy, B. P., Gordon, S., Henders, A. K. Nyholt, D. R., … Visscher, P. M. (2010). Common SNPs explain a large proportion of the heritability for human height. Nature genetics, 42(7), 565–9. doi:10.1038/ng.608

Yang, J., Manolio, T. a, Pasquale, L. R., Boerwinkle, E., Caporaso, N., Cunningham, J. M., … Visscher, P. M. (2011). Genome partitioning of genetic variation for complex traits using common SNPs. Nature genetics, 43(6), 519–25. doi:10.1038/ng.823

Zhang, Y., Brady, M., & Smith, S. (2001). Segmentation of brain MR images through a hidden Markov random field model and the expectation-maximization algorithm. IEEE Transactions on Medical Imaging, 20(1), 45–57. doi:10.1109/42.906424

